# scRNA-seq reveals ACE2 and TMPRSS2 expression in TROP2^+^ Liver Progenitor Cells: Implications in COVID-19 associated Liver Dysfunction

**DOI:** 10.1101/2020.03.23.002832

**Authors:** Justine Jia Wen Seow, Rhea Pai, Archita Mishra, Edwin Shepherdson, Tony Kiat Hon Lim, Brian KP Goh, Jerry KY Chan, Pierce KH Chow, Florent Ginhoux, Ramanuj DasGupta, Ankur Sharma

## Abstract

The recent pandemic of coronavirus disease 2019 (COVID-19) is caused by severe acute respiratory syndrome coronavirus 2 (SARS-CoV-2). COVID-19 was first reported in China (December 2019) and now prevalent in ∼170 countries across the globe. Entry of SARS-CoV-2 into mammalian cells require the binding of viral Spike (S) proteins to the ACE2 (angiotensin converting enzyme 2) receptor. Once entered the S protein is primed by a specialised serine protease, TMPRSS2 (Transmembrane Serine Protease 2) in the host cell. Importantly, beside respiratory symptoms, consistent with other common respiratory virus infection when patients become viraemic, a significant number of COVID-19 patients also develop liver comorbidities. We explored if specific target cell-type in the mammalian liver, could be implicated in disease pathophysiology other than the general deleterious response to cytokine storms. Here we employed single-cell RNA-seq (scRNA-seq) to survey the human liver and identified potentially implicated liver cell-type for viral ingress. We report the co-expression of ACE2 and TMPRSS2 in a TROP2^+^ liver progenitor population. Importantly, we fail to detect the expression of ACE2 in hepatocyte or any other liver (immune and stromal) cell types. These results indicated that in COVID-19 associated liver dysfunction and cell death, viral infection of TROP2^+^ progenitors in liver may significantly impaired liver regeneration and could lead to pathology.

**Highlights:**

- EPCAM^+^ Liver progenitors co-express ACE2 and TMPRSS2
- ACE2 and TMPRSS2 expression is highest in TROP2^high^ progenitors
- ACE2 and TMPRSS2 cells express cholangiocyte biased fate markers
- ACE2 and TMPRSS2 positive cells are absent in human fetal liver

## Introduction

Since December of 2019, the SARS-CoV-2 pandemic has impacted millions of lives worldwide. As of 23^rd^ March 2020, more than 340,000 people are reported to be infected, with ∼4-5% mortalities (https://coronavirus.jhu.edu/map.html). The SARS-CoV-2 is a single-stranded RNA virus belonging to the Coronaviridae family of zoonotic viruses that infect mammals and birds (Andersen et al., 2020). The novel SARS-CoV-2 was first isolated from lung airway epithelial cells of patient with pneumonia (Chen et al., 2020). Since then it has been reported that SARS-CoV-2 employ receptor angiotensin converting enzyme 2 (ACE2) for entry into human cells and utilise Transmembrane Serine Protease 2 (TMPRSS2) for Spike (S) Protein priming (Hoffmann et al., 2020). SARS-CoV-2 shares ∼80% sequence similarity with SARS-CoV and ∼50% with Middle East respiratory syndrome coronavirus (MERS-CoV), all of which cause severe respiratory symptoms (Hoffmann et al., 2020). Moreover, in addition to respiratory disease, SARS and MERS are known to cause liver impairments (Alsaad et al., 2017; Chau et al., 2004; Zhang et al., 2020).

Importantly, SARS-CoV-2 RNA was discovered in stool sample of the first US patient, indicating gastrointestinal tract infection (Holshue et al., 2020). A recent study reported 14-53% cases with higher levels of alanine aminotransferase (ALT) and aspartate aminotransferase (AST) in the liver of COVID-19 patients (Huang et al., 2020; Zhang et al., 2020). Moreover, these symptoms were elevated in intensive care unit (ICU) patients compared to patients who did not require ICU (Huang et al., 2020). It remains to be investigated whether SARS-CoV-2 directly infect liver cells. In addition, concerns have been raised on effect of SARS-CoV-2 infection on pre-existing liver condition (Gu et al., 2020; Mao et al., 2020; Zhang et al., 2020). Since SARS-CoV-2 binds to ACE2 and require TMPRSS2 for activation, we surveyed the human liver (from tumor and adjacent normal regions of hepatocellular carcinoma patients) by sc-RNA-seq to identify which cell type co-express these two genes.

Here we report that ACE2 and TMPRSS2 are co-expressed in only one sub-population in human liver. Based on expression of cell type specific markers ALB (Albumin), KRT (Keratin), EPCAM and unique expression pattern of TROP2 (TACSTD2) and SOX9 (SRY-box 9), we annotated this population as liver progenitors. Our results suggest SARS-CoV-2 binding receptor ACE2 is only expressed on TROP2^high^ cholangiocyte-biased progenitors while serine protease TMPRSS2 is expressed by TROP2^high^ and TROP2^int^ population. These results indicate that SARS-CoV-2 infection might preferentially infect the TROP2^high^ cholangiocyte-biased progenitor pool, thereby compromising the regenerative abilities of infected liver and/or contributing to liver pathology.

## Results

### Expression of ACE2 and TMPRSS2 in human liver scRNA-seq atlas

We performed scRNA-seq on human liver tissue obtained from tumor and adjacent normal tissue of hepatocellular carcinoma patients (Sharma et al., in revision). In total, we analysed ∼74,000 cells and employed Louvain algorithm for clustering of these cells. We identified 29 clusters based on the expression of cell type specific genes. These clusters were further annotated into hepatocyte, epithelial progenitors, lymphoid and myeloid cells (Figure 1A) and observed specific enrichment of EPCAM^+^ and SOX9^+^ epithelial progenitor cluster in normal liver (Figure 1B). We identified the specific markers for every cluster (Figure 1C) and observed the co-expression of KRT18 and KRT19 in epithelial progenitor cluster. Next, we investigated which cell types in the human liver co-express SARS-CoV-2 binding receptor ACE2 and the priming enzyme TMPRSS2. Our analysis revealed the specific expression of ACE2 and TMPRSS2 in the epithelial progenitor cluster (Figure 1D and 1E). This suggests that EPCAM^+^ liver progenitors express machinery for both SARS-CoV-2 entry (ACE2) as well as priming (TMPRSS2) and might be susceptible for viral infection leading to liver dysfunction.

**Figure-1.**
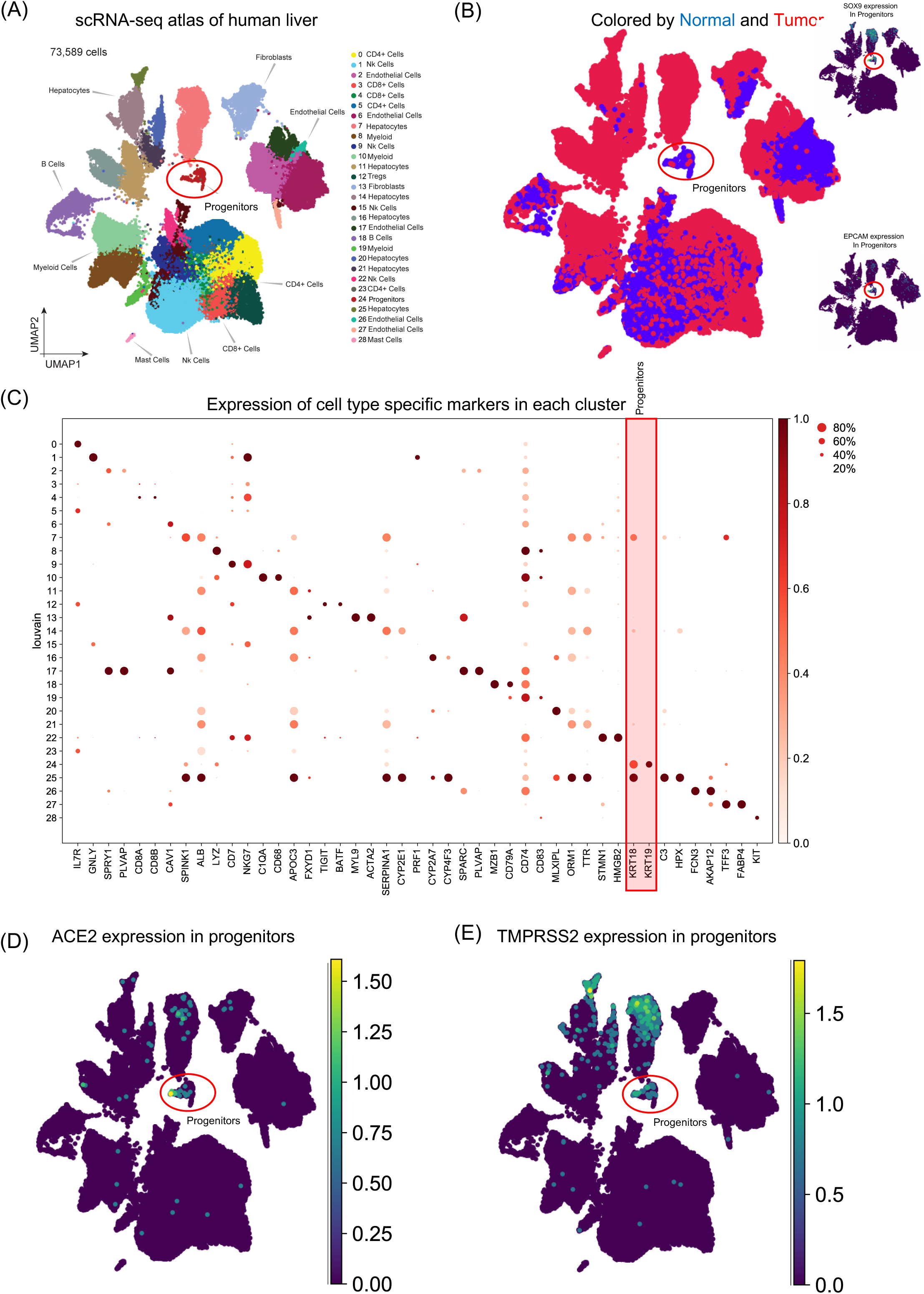
Expression of ACE2 and TMPRSS2 in human liver. (A) Louvain clustering of ∼74,000 sc-RNA-seq libraries from 14 HCC patients identifies 29 clusters in human liver. (B) Louvain clusters colored by normal (blue) and tumor (red) cell types. Note the UMAP expression of SOX9 and EPCAM in progenitor cluster. (C) Dot-plot depicting cell type specific markers in human liver. Note the co-expression of KRT18 and KRT19 in progenitor cluster indicating presence of bi-potent cells in this cluster. Expression of SARS-CoV-2 entry receptor (D) ACE2 and S-protein converting enzyme (E) TMPRSS2 in human liver progenitor cluster.

### TROP2+ liver epithelial progenitors express ACE2 and TMPRSS2

A recent scRNA-seq study has suggested the heterogeneity in EPCAM^+^ liver epithelial progenitors (Aizarani et al., 2019). Therefore, we further sub-clustered the epithelial cells (hepatocytes and progenitors) to understand the nature of ACE2 expressing liver progenitors (Figure 2A). Sub-clustering also showed the predominant presence of normal liver cells in the progenitor cluster (Figure 2B). Further we detected the presence of these cells in adjacent normal region from multiple individuals (Figure 2C). Interestingly, lower abundance of these cells in human liver is in concordance with the rare stem-like or progenitor population in epithelial tissues. Next, we analysed the expression of hepatocyte, cholangiocyte and bi-potent markers in these clusters. The progenitor cluster specifically expressed EPCAM (progenitor marker) as well as KRT19 and CFTR (Cystic fibrosis transmembrane conductance regulator) which are known to be expressed in progenitors with a cholangiocyte fate bias (Figure 2D) (Cohn et al., 1993). Importantly, we failed to detect expression of hepatocyte fate bias genes, ASGR1 (Asialoglycoprotein receptor 1) and ALB in this cluster. Since this progenitor cluster demonstrated bias for cholangiocyte fate we further investigated the expression of TROP2 gene. The TROP2 expression is known to mark the fate of liver epithelial progenitors where lower TROP2 expression is linked with hepatocyte fate and TROP2^high^ cells with cholangiocyte fate (Aizarani et al., 2019).

**Figure-2.**
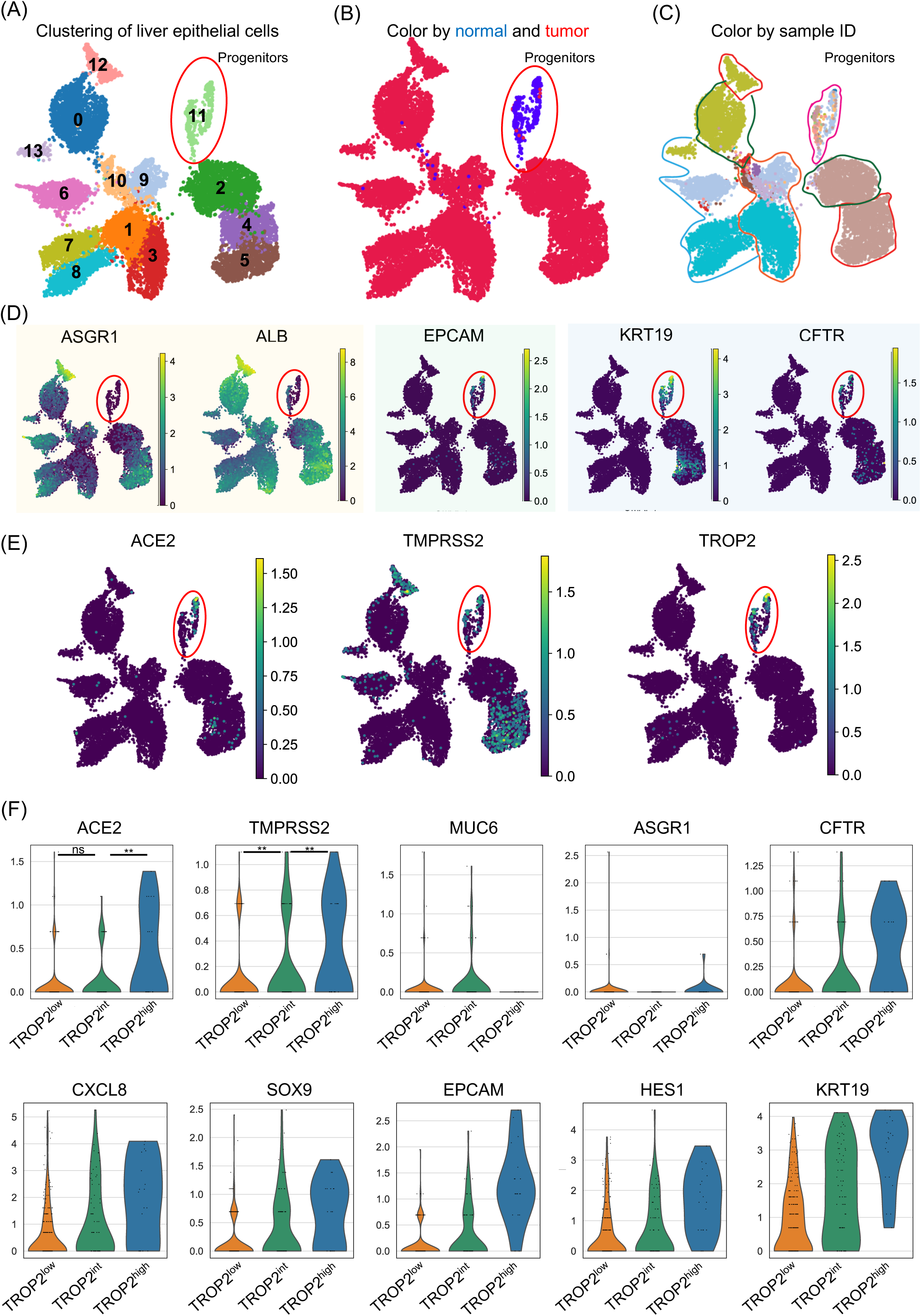
Expression of ACE2 and TMPRSS2 in TROP2^+^ liver progenitors. (A) Louvain clustering of ∼10,000 sc-RNA-seq libraries from epithelial cells identifies 12 sub-clusters in human liver. (B) Louvain clusters colored by normal (blue) and tumor (red) cell types. Note the predominate normal cells in progenitor cluster. (C) Louvain clusters colored by sample ID note the representation of multiple sample IDs in progenitor cluster. (D) Expression of hepatocyte fate biased (ASGR1 and ALB), bi-potent (EPCAM) and cholangiocyte fate biased (KRT19 and CFTR) genes in progenitor cluster (circled red). (E) Expression of ACE2, TMPRSS2 and TROP2 in progenitor cluster (circled red). (F) Expression of ACE2 and TMPRSS2 in TROP2^low^ (orange), TROP2^int^ (green) and TROP2^high^ (blue) cells (note the highest expression of ACE2 and TMPRSS2 in TROP2^high^ cells as well as expression of other cell fate markers). (The progenitor cluster was binned into TROP2^low^ (orange), TROP2^int^ (green) and TROP2^high^ (blue) cells)

Recently, Aizarani et al., demonstrated the progenitor-like properties of TROP2^+^ cells where TROP2^Int^ cells demonstrated highest organoid forming efficiency followed by TROP2^high^ cells, while TROP2^low^ cells failed to generate organoids (Aizarani et al., 2019). Therefore, we investigated whether any of the epithelial clusters co-expressed ACE2, TMPRSS2, and TROP2. Notably, we observed that only the EPCAM^+^ progenitor cluster expressed all three genes (Figure 2E). Next we sub-divided this cluster into TROP2 low, intermediate and high cells and investigated the expression of ACE2, TMPRSS2 and other cell fate markers. Remarkably, we observed that TROP2^high^ cells expressed the highest levels of ACE2 and TMPRSS2, followed by TROP2^Int^ and TROP2^low^ cells (Figure 2F). Our analysis revealed that TROP2^Int^ (bi-potent) cells also express MUC6 and SOX9, whereas TROP2^high^ (cholangiocyte fate bias) cells express makers such as CFTR, CXCL8, HES1 and KRT19. Taken together, our results suggest that SARS-CoV-2 can infect TROP2^high^ cells via ACE2 and TMPRSS2, thereby contributing to liver dysfunction by compromising the ability of the human liver to regenerate cholangiocytes.

Additionally given the recent discussion on possible intrauterine vertical transmission of SARS-CoV-2, we also analysed human fetal liver scRNA-seq atlas at 16 and 21 week EGA. Our analysis indicated very low to negligible level of ACE2 and an absence of TMPRSS2 expression in EPCAM+ fetal liver cells (Figure 3A-D). Our limited analysis suggest that fetal liver may not express the machinery required for SARS-CoV-2 infection.

**Figure-3.**
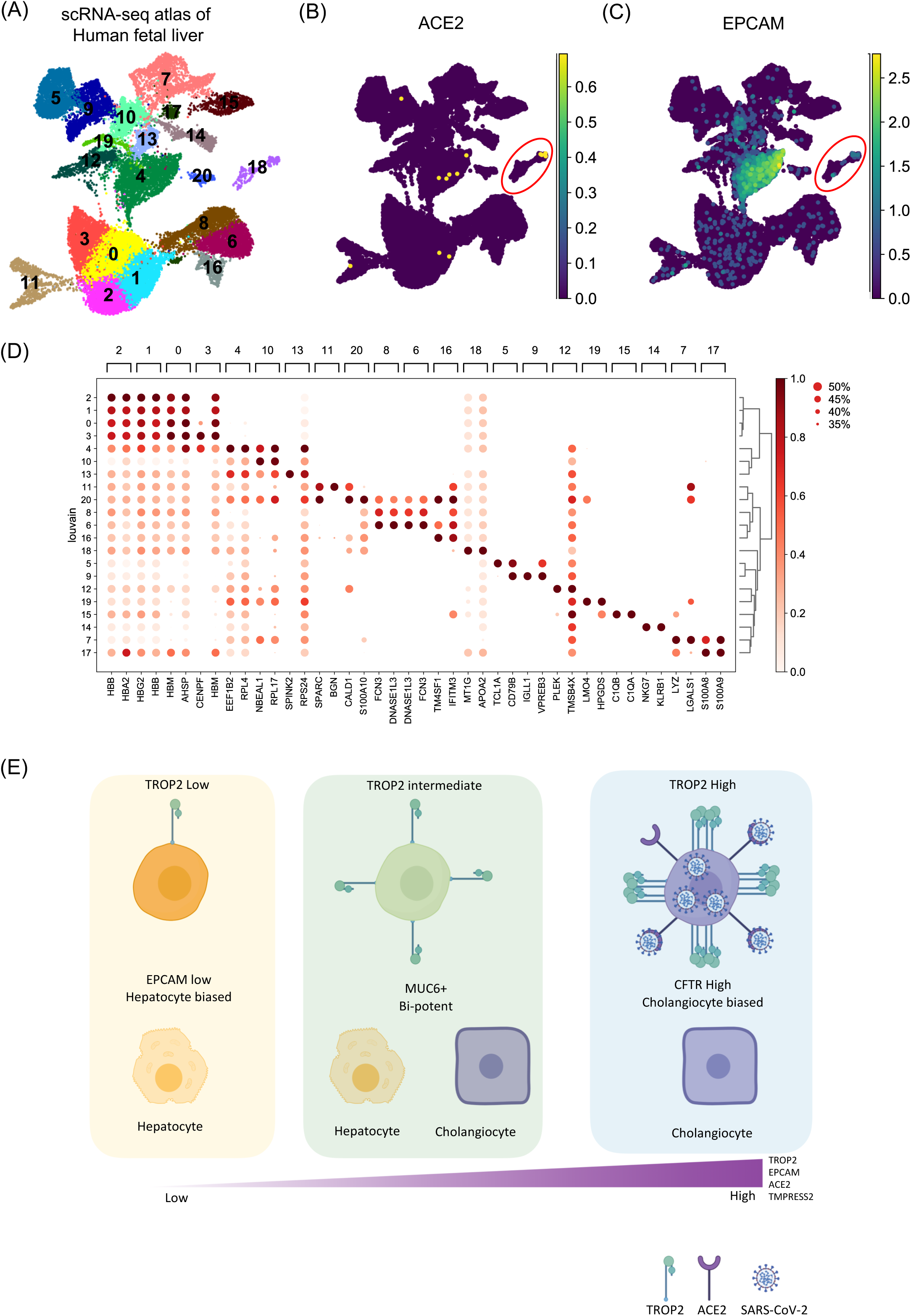
ACE2 expression in human fetal liver. (A) Louvain clustering of ∼35,000 sc-RNA-seq libraries from human fetal liver. Expression of (B) ACE2 and (C) EPCAM. Note the very low expression of ACE2 in EPCAM^+^ human fetal liver cells (circled red). (D) Dot-plot depicting cell type specific markers in human fetal liver.

## Discussion

In recent reports of SARS-CoV-2 pandemic in human population, the presence of viral mRNA in infected patient’s stool suggests a potential GI (gastrointestinal) tract infection in COVID-19 patients. SARS-CoV-2 can reach the liver either through the general circulation once patient has become viraemic or through transmigration through the GI tract. We surveyed human liver scRNA-seq data to understand the expression pattern of ACE2 and TMPRSS2 gene which are essential for SARS-CoV-2 entry into human cells. Our analysis reveals that in human liver only EPCAM^+^ progenitors co-express genes for viral entry (ACE2) and S-protein priming (TMPRSS2). Further analyses revealed the specific expression of ACE2 and TMPRSS2 in TROP2^high^ cells. These results indicate that ACE2 and TMPRSS2 are specifically present in liver progenitors with a cholangiocyte fate bias, suggesting SARS-CoV-2 may be affecting cholangiocyte precursors, thereby potentially impeding the homeostasis of the cholangiocyte pool. Recent studies have reported the expression of ACE2 in cholangiocytes, however, did not reflect over the heterogeneity of ACE2^+^ population (Chai et al., 2020). In this study we explored the heterogeneity of ACE2^+^ cells and systematically characterised ACE2 and TMPRSS2 co-expression as hallmarks of TROP2^+^ epithelial progenitors.

Our study reveals the potential of SARS-CoV-2 to infect TROP2^+^ progenitor like cells in human liver. It is important to note that TROP2 is expressed in multiple epithelial progenitors (Crowell et al., 2019; Okabe et al., 2009; Vallone et al., 2016). In the future it will be important to survey other GI tract tissues at single-cell level for the expression of ACE2 and TMPRSS2 and their associated transcriptomes. Given the GI tract infection and multi-organ failure in COVID-19, it is important to understand whether other progenitor-like cells are also susceptible to SARS-CoV-2 infection. Moreover, TROP2 expression has been associated with amplifying progenitor cells in partial hepatectomy mouse model (Okabe et al., 2009) indicating the important role of TROP2^+^ cells in liver regeneration. Taken together, our analysis suggests that adult human liver TROP2^+^ progenitors could be a prime target of SARS-CoV-2 (Figure 3E).

General hepatocyte cell damage from cytokine storms in ill patients with viraemia from respiratory viral infection is not uncommon. Such hepatocyte damage are usually transient and the resulting liver regeneration usually restores liver function efficiently. In the case of COVID-19, however the predilection of the SARS-CoV-2 virus for cholangiocyte precursor cells may significantly impair liver regeneration. Clinicians looking after patients with COVID-19 should be alerted to the possibility of progressive liver deterioration in patients with serious SARS-Cov-2 viraemia. This study demonstrates the power of single-cell RNA-seq to understand the pathobiology of COVID-19 and pave the way for similar studies to understand the effect of SARS-CoV-2 on different tissue and cell types.

## Figure legends

**Figure-4 Heterogeneity of ACE2 and TMPRSS2 expression in liver progenitors**

Schematic of TROP2 expression level and cell fate choices in adult human liver progenitor cells. TROP2^high^ cells express genes exhibiting cholangiocyte fate bias. The TROP2^high^ cells also express higher level of ACE2 and TMPRESS2 making these cells more susceptible for SARS-CoV-2 infection, indicating implications in COVID-19 associated liver dysfunctions.

## Experimental Methods

### Tissue acquisition

The fresh tissue samples were obtained from Singapore General Hospital (SGH) and National University (NUH) with written consent and approval from central institution review board of singhealth (CIRB2012/669/B) for study on liver cancer. The samples were delivered in with MACS Tissue Storage Solution (Miltenyi, Cat#:130-100-008).

### Human fetal liver samples

The donation of fetal liver tissues for research was approved by the Centralised Institutional Research Board of the Singapore Health Services in Singapore followed by proper international ethical guidelines and in accordance with a favorable ethical opinion from Singapore SingHealth and National Health Care Group Research Ethics Committees. Women gave written informed consent for the donation of fetal tissue to research nurses who were not directly involved in the research, or in the clinical treatments of women participating in the study, as per the Polkinghorne guidelines. This protocol was reviewed on an annual basis by the Centralised Institutional Research Board (IRB2013/837/D), including annual monitoring of any adverse events, for which there had been none. All fetal liver tissues were obtained from 2nd trimester (16 and 21 weeks EGA) elective pregnancy terminations carried out for socio-psychological reasons. All fetuses were considered structurally normal on ultrasound examination prior to termination and by gross morphological examination following termination. In total 2 fetuses of 16 and 21 weeks EGA were used for this study.

### Tissue processing

Tissues were transferred immediately using the following procedure. Tissue was transferred to a sterile 10mm^2^ tissue culture dish and cut into very small fragments. The dissociation buffer consisted of 0.43mg/ml of Collagenase IV (Thermofisher, Cat#: 17104019) and 0.172mg/ul of DNAse1 (Worthington, Cat#: LS002147) dissolved in PBS (Thermofisher, Cat#: 20012-043). The tissue was digested in dissociation buffer for 30-40 minutes depending on sample size at 37°C with constant shaking at 220rpm while keeping the falcon tube in a slanted position. The solution was re-suspended with 10ml pipette followed by 18g needle. The 1% BSA PBS solution was added to the digested tissue and then the solution was passed through a 70um filter before centrifuging at 800g for 6 minutes at 4°C. Cells were treated with 5ml of 1x RBC lysis buffer (Biolegend, Cat#: 420301) on ice for 10-15 minutes. 1% BSA PBS solution was then added and the cells were passed through 40um filter. Cells were dissolved in 1% BSA PBS solution prior to counting.

### Data processing using Cell Ranger software

Sequenced fastq files are aligned, filtered, barcoded and UMI counted using Cell Ranger Chromium Single Cell RNA-seq version 2.0.2, by 10X Genomics with Cell Ranger, GRCh38 database (version 1.2.0) as the human genome reference. All 62 sectors are aggregated using *cellranger aggr* by normalizing all runs to the same sequencing depth.

### Clustering and downstream analysis

Downstream analysis was done using Scanpy, a scalable Python-based package (version 1.4) designed for single cell gene expression datasets. Scanpy implements numerous functions from preprocessing to visualization, clustering, differential gene expression, and trajectory inference analysis on Jupyter Notebooks. Parameters used in each function are manually curated to portray the best clustering of cells. In preprocessing, cells are filtered based on the criteria of expressing a minimum of 200 genes and a gene which is expressed by a minimum of 30 cells. Dying cells with a mitochondrial percentage of more than 5% are excluded. Cell count was normalized using *scanpy*.*api*.*pp*.*normalize_per_cell* with a scaling factor of 10,000 whereas gene expression was scaled to unit variance and mean value of 0 using *scanpy*.*api*.*pp*.*scale*. Dimension reduction starts with PCA using *scanpy*.*api*.*tl*.*pca*; the number of PCs used in each clustering exercise varies depending on the importance of embeddings to be included. In the interest of crisp clustering, we first calculated neighborhood graph (*scanpy*.*api*.*pp*.*neighbors*) of cells. Best matched k-Nearest Neighbor is automatically weighted by the algorithm to compute the best UMAP topology (*scanpy*.*api*.*tl*.*umap*, minimum distance between 0.3 to 0.5) which is consistently used throughout this paper. Louvain method (*scanpy*.*api*.*tl*.*louvain*) is then used to detect a community of similar cells. By default, Louvain’s resolution parameter is set to the maximum value of 1.0, this in theory finds more and smaller clusters. In our experiments, the value is set between 0.6 to 1. Genes are then ranked using *scanpy*.*api*.*tl*.*rank_genes_groups* (Benjamini Hochberg, t-test overestimated variance with adjusted p-value). Cell types was manually and iteratively assigned based on overlaps of literature curated and statistically ranked genes. To leverage the heterogeneity of this dataset, we used partition-based graph abstraction (PAGA, *scanpy*.*api*.*tl*.*paga*) to reconstruct lineage between cell types. This lineage trajectory provides a continuous cell type transition from the assigned discrete cell types. The thickness of the edges represents connectivity scores, an entropy-based measure provided by PAGA indicating relatedness between clusters; spurious connections are discarded while tuning thresholds.

## Acknowledgments

We thank all patients and families involved in this study. We thank members of GIS, SIgN, SGH, NCCS and KKH teams and Liver TCR group for useful discussions. We thank Sin Chi Chew, the GIS sequencing core and the SIgN FACS core platforms for their help and support. This work is supported by National Medical Research Council (NMRC, Singapore) grant TCR15Jun006, Agency for Science, Technology and Research (A*STAR) core funds to R.D.G. and F.G.. A.S. is supported by NMRC young investigator grant (OFYIRF18nov-0056).

## Author Contributions

Conceptualization: A.S., P.K.H.C., F.G., and R.D.G.; Methodology: A.S., J.J.W.S.,; Investigation: A.S., J.J.W.S., A.M.,; Formal analysis; A.S., J.J.W.S.,; Data Curation: A.S., J.J.W.S.; Writing-original draft A.S.; Writing-review and editing P.K.H.C., F.G., and R.D.G.; Funding Acquisition; P.K.H.C., F.G., A.S. and R.D.G; Resources, J.C., T.K.H.L., B.K.P.H, P.K.H.C., F.G., and R.D.G.; Supervision, A.S.

## Declaration of Interests

No conflict of interest to declare

## Notes

https://coronavirus.jhu.edu/map.html

## References

Aizarani, N., Saviano, A., Sagar, Mailly L., Durand, S., Herman, J.S., Pessaux, P., Baumert, T.F., and Grün, D. (2019). A human liver cell atlas reveals heterogeneity and epithelial progenitors. Nature 572, 199–204.

Alsaad, K.O., Hajeer, A.H., Balwi, M.A., Moaiqel, M.A., Oudah, N.A., Ajlan, A.A., AlJohani, S., Alsolamy, S., Gmati, G.E., Balkhy, H., et al. (2017). Histopathology of Middle East respiratory syndrome coronovirus (MERS-CoV) infection - clinicopathological and ultrastructural study. Histopathology 72, 516–524.

Andersen, K.G., Rambaut, A., Lipkin, W.I., Holmes, E.C., and Garry, R.F. (2020). The proximal origin of SARS-CoV-2. Nat Med 1–3.

Chai, X., Hu, L., Zhang, Y., Han, W., Lu, Z., Ke, A., Zhou, J., Shi, G., Fang, N., Fan, J., et al. (2020). Specific ACE2 Expression in Cholangiocytes May Cause Liver Damage After 2019-nCoV Infection. Biorxiv 2020.02.03.931766.

Chau, T.-N., Lee, K.-C., Yao, H., Tsang, T.-Y., Chow, T.-C., Yeung, Y.-C., Choi, K.-W., Tso, Y.-K., Lau, T., Lai, S.-T., et al. (2004). SARS-associated viral hepatitis caused by a novel coronavirus: Report of three cases. Hepatology 39, 302–310.

Chen, N., Zhou, M., Dong, X., Qu, J., Gong, F., Han, Y., Qiu, Y., Wang, J., Liu, Y., Wei, Y., et al. (2020). Epidemiological and clinical characteristics of 99 cases of 2019 novel coronavirus pneumonia in Wuhan, China: a descriptive study. Lancet Lond Engl 395, 507–513.

Cohn, J.A., Strong, T.V., Picciotto, M.R., Nairn, A.C., Collins, F.S., and Fitz, J.G. (1993). Localization of the cystic fibrosis transmembrane conductance regulator in human bile duct epithelial cells. Gastroenterology 105, 1857–1864.

Crowell, P.D., Fox, J.J., Hashimoto, T., Diaz, J.A., Navarro, H.I., Henry, G.H., Feldmar, B.A., Lowe, M.G., Garcia, A.J., Wu, Y.E., et al. (2019). Expansion of Luminal Progenitor Cells in the Aging Mouse and Human Prostate. Cell Reports 28, 1499-1510.e6.

Gu, J., Han, B., and Wang, J. (2020). COVID-19: Gastrointestinal manifestations and potential fecal-oral transmission. Gastroenterology.

Hoffmann, M., Kleine-Weber, H., Schroeder, S., Krüger, N., Herrler, T., Erichsen, S., Schiergens, T.S., Herrler, G., Wu, N.-H., Nitsche, A., et al. (2020). SARS-CoV-2 Cell Entry Depends on ACE2 and TMPRSS2 and Is Blocked by a Clinically Proven Protease Inhibitor. Cell.

Holshue, M.L., DeBolt, C., Lindquist, S., Lofy, K.H., Wiesman, J., Bruce, H., Spitters, C., Ericson, K., Wilkerson, S., Tural, A., et al. (2020). First Case of 2019 Novel Coronavirus in the United States. New Engl J Medicine 382, 929–936.

Huang, C., Wang, Y., Li, X., Ren, L., Zhao, J., Hu, Y., Zhang, L., Fan, G., Xu, J., Gu, X., et al. (2020). Clinical features of patients infected with 2019 novel coronavirus in Wuhan, China. Lancet Lond Engl 395, 497–506.

Mao, R., Liang, J., Shen, J., Ghosh, S., Zhu, L.-R., Yang, H., Wu, K.-C., Chen, M.-H., Union, C.S. of I., Chinese Elite IBD, and Committee, C.I.Q.C.E.C. (2020). Implications of COVID-19 for patients with pre-existing digestive diseases. Lancet Gastroenterology Hepatology.

Okabe, M., Tsukahara, Y., Tanaka, M., Suzuki, K., Saito, S., Kamiya, Y., Tsujimura, T., Nakamura, K., and Miyajima, A. (2009). Potential hepatic stem cells reside in EpCAM+ cells of normal and injured mouse liver. Dev Camb Engl 136, 1951–1960.

Vallone, V.F., Leprovots, M., Strollo, S., Vasile, G., Lefort, A., Libert, F., Vassart, G., and Garcia, M.-I. (2016). Trop2 marks transient gastric fetal epithelium and adult regenerating cells after epithelial damage. Dev Camb Engl 143, 1452–1463.

Zhang, C., Shi, L., and Wang, F.-S. (2020). Liver injury in COVID-19: management and challenges. Lancet Gastroenterology Hepatology.

